# Disentangling evoked and induced activity from single-trial neurophysiological time series

**DOI:** 10.64898/2026.10.19.760024

**Authors:** Oskari Ahola, Lisa Haxel, Ulf Ziemann, Risto J. Ilmoniemi, Pantelis Lioumis

**Affiliations:** Department of Neuroscience and Biomedical Engineering, Aalto University, Espoo, Finland; Translational Neurotechnology Laboratory, Department of Neurosurgery, TUM Klinikum Rechts der Isar, Technical University of Munich, Germany; Institute of Computational Biology, Helmholtz Munich, Neuherberg, Germany; Department of Neurology and Stroke, University of Tübingen, Tübingen, Germany; Hertie Institute for Clinical Brain Research, University of Tübingen, Tübingen, Germany; BioMag Laboratory, HUS Medical Imaging Center; Aalto University; University of Helsinki; Helsinki University Hospital, Helsinki, Finland; Department of Psychology and Logopedics, University of Helsinki, Helsinki, Finland

## Abstract

Evoked and induced responses are fundamental neurophysiological readouts in the development and assessment of treatments for disorders with impaired neural function. Distinguishing these responses is essential for interpreting single-trial variability, identifying the determinants of these responses, and enabling real-time probing of cortical responsiveness. However, because of dynamic overlap, separating evoked and induced activity remains unsolved at the single-trial level when instantaneous fluctuations of cortical responsiveness are of interest.

We developed the Evoked–Ongoing/Induced Separator (EOIS), a framework for separating event-evoked, ongoing, and event-modulated induced activity from single-trial data. EOIS recovers evoked and ongoing/induced components by leveraging their distinct phase-consistency features in MEG and real-time EEG, enabling subject-agnostic single-trial time-series analysis and attribution of cortical response variability to specific components. By delivering component-resolved time series, EOIS enables biomarker refinement, pharmacodynamic readouts, and closed-loop neuromodulation, facilitating translational deployment across individuals and care settings.

## 1 Main

Evoked responses are components of cortical activity that are time- and phase-locked to an event. In trial-averaged data, these components are clearly observable because non-phase-locked activity and noise converge toward zero over trials. However, at the single-trial level, evoked responses are dynamically intermixed with ongoing and event-modulated (induced) non-phase-locked activity, which can confound inferences drawn from either measure.

Single-trial electroencephalographic and magnetoencephalographic (EEG and MEG) readouts provide biomarkers for cortical and corticospinal responsiveness during different brain states. These readouts have been used to investigate the relationships between ongoing activity and stimulation reactivity^1–4^, identify brain states associated with high or low excitability^5^, guide spatial cortical response mapping and thresholding procedures for transcranial magnetic stimulation (TMS)^6,7^, and support brain–computer interfaces (BCIs)^8,9^. When investigating the dynamics of single-trial responses, the ongoing and stimulation-modulated activity should be accounted for because these components are not independent. Without temporal component separation, pre-stimulus dynamics and ongoing background activity are present in the post-stimulus data, which confound attributions of post-stimulus variability.

Existing approaches use constrained filters to estimate single-trial event-related subcomponent amplitudes and latencies^5,10,11^. However, they require subject-specific calibration; robust, generalizable methods for distinguishing single-trial evoked and ongoing/induced activity without subject-specific priors have yet to be demonstrated. We address this by enabling real-time estimation of evoked and ongoing/induced single-trial time series without subject-specific calibration, enhancing generalizability across individuals and sessions.

We developed the Evoked–Ongoing/Induced Separator (EOIS), a temporal signal separation algorithm. EOIS leverages convergent features of loosely phase-locked and non-phase-locked activity to separate evoked and ongoing/induced components at the single-trial level in MEG and EEG. The method is applicable to multiple event-based data modalities, is subject-agnostic, and is suitable for real-time EEG applications.

**Fig 1:**
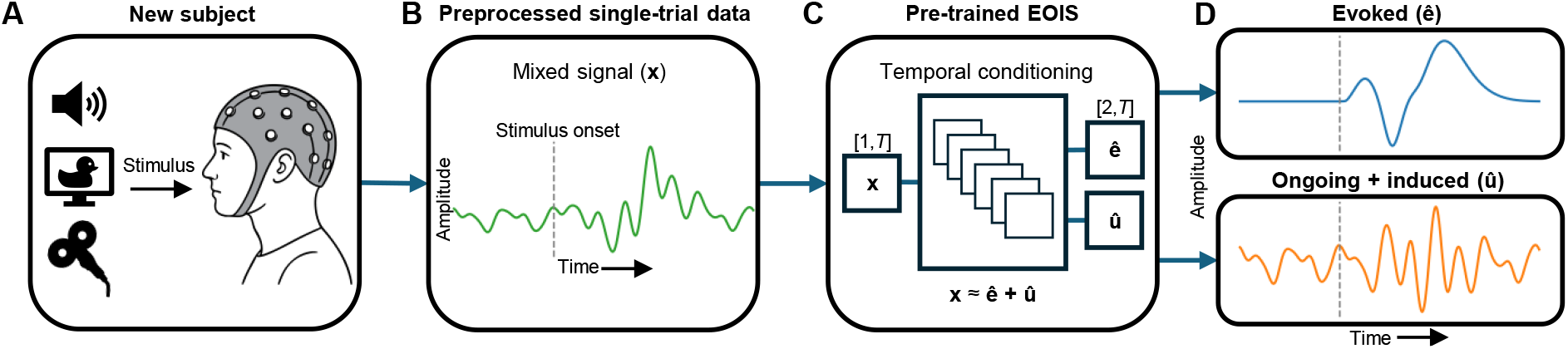
Framework for separating evoked and ongoing/induced activity from single-trial neurophysiological time series.**A**: A new subject is given a stimulus, *e*.*g*., auditory, visual, or TMS. **B**: Each EEG/MEG trial includes ongoing non-phase-locked activity before the stimulus, post-stimulus non-phase-locked (induced) activity, and post-stimulus phase-locked (evoked) activity. **C**: A single-trial time series of length *T* is provided to EOIS, which separates the input signal into evoked and on-going/induced components (**D**) using dilated residual blocks^12–15^.

## 2 Results

### 2.1 Amplitude and frequency representations

We evaluated EOIS on visual (VIS) and auditory (AUD) MEG and on TMS–EEG, with and without across-trial acausal filter calibration (TMS and TMS-st). Across-trial phase locking was quantified using pairwise phase consistency (PPC)^16^ computed from Morlet wavelet transforms for both evoked and ongoing/induced components. EOIS consistently recovers these components in the time–amplitude and time–frequency domains across leave-one-subject-out (LOSO) folds. Table 1 and Fig. 2 summarize results by train/test data modality pair within stimulation-relevant windows. Evoked components showed strong cross-subject and trial-wise consistency in both domains. Ongoing/induced components, which may be time-locked to stimulus onset but are not phase-locked, were more dispersed in the time–frequency domain and averaged to near zero in absolute amplitude. Accordingly, ongoing/induced activity yielded near-zero PPCs, whereas evoked activity showed elevated PPCs without approaching an unrealistic limit of 1, which would imply no temporal jitter.

**Table 1:** Recovery and phase-consistency metrics for unseparated, ongoing/induced, and evoked components across train/test modality pairs (LOSO). Values are median *±* median absolute deviation (MAD) across subjects. PPC quantifies post-stimulus phase locking from 0 to 400 ms of post-stimulus data (EEG: 4–40; MEG: 4–30; low 4–15; high 15–30 Hz); *R*^2^ Trial and *R*^2^ Average evoked quantify reconstruction accuracy for single-trial activity and across-trial averaged evoked activity for the post-stimulus data. Visual (VIS), auditory (AUD), TMS, and causally across trials processed TMS (TMS-st) denote the train/test modalities. *N*_subjects_ is the number of test subjects. These are also the number of training/validation subjects (excluding the left out subject) when the trained model type is equal to the tested data type (see Section 5.2).

| Metric | VIS/VIS | VIS/AUD | AUD/VIS | AUD/AUD | TMS/TMS | TMS-st/TMS-st |
| --- | --- | --- | --- | --- | --- | --- |
| $N_{\text{subjects}}$ | 219 | 49 | 49 | 92 | 52 | 52 |
| PPC(Unseparated)<sub>full</sub> | $0.230 \pm 0.048$ | $0.220 \pm 0.032$ | $0.235 \pm 0.042$ | $0.211 \pm 0.040$ | $0.139 \pm 0.060$ | $0.178 \pm 0.062$ |
| PPC(Ongoing/Induced)<sub>full</sub> | $0.015 \pm 0.008$ | $0.019 \pm 0.016$ | $0.061 \pm 0.023$ | $0.014 \pm 0.014$ | $0.023 \pm 0.012$ | $0.024 \pm 0.011$ |
| PPC(Evoked)<sub>full</sub> | $0.736 \pm 0.044$ | $0.716 \pm 0.047$ | $0.760 \pm 0.043$ | $0.796 \pm 0.034$ | $0.663 \pm 0.066$ | $0.706 \pm 0.066$ |
| PPC(Unseparated)<sub>low</sub> | $0.414 \pm 0.080$ | $0.434 \pm 0.063$ | $0.419 \pm 0.081$ | $0.410 \pm 0.074$ | $0.260 \pm 0.126$ | $0.309 \pm 0.117$ |
| PPC(Ongoing/Induced)<sub>low</sub> | $0.014 \pm 0.010$ | $0.023 \pm 0.031$ | $0.066 \pm 0.024$ | $0.015 \pm 0.019$ | $0.028 \pm 0.015$ | $0.034 \pm 0.019$ |
| PPC(Evoked)<sub>low</sub> | $0.683 \pm 0.072$ | $0.655 \pm 0.074$ | $0.667 \pm 0.072$ | $0.699 \pm 0.064$ | $0.422 \pm 0.121$ | $0.511 \pm 0.111$ |
| PPC(Unseparated)<sub>high</sub> | $0.070 \pm 0.033$ | $0.029 \pm 0.029$ | $0.073 \pm 0.032$ | $0.034 \pm 0.026$ | $0.096 \pm 0.053$ | $0.127 \pm 0.048$ |
| PPC(Ongoing/Induced)<sub>high</sub> | $0.014 \pm 0.009$ | $0.016 \pm 0.017$ | $0.057 \pm 0.024$ | $0.008 \pm 0.020$ | $0.022 \pm 0.013$ | $0.024 \pm 0.013$ |
| PPC(Evoked)<sub>high</sub> | $0.795 \pm 0.034$ | $0.778 \pm 0.036$ | $0.838 \pm 0.028$ | $0.863 \pm 0.024$ | $0.728 \pm 0.047$ | $0.767 \pm 0.058$ |
| $R^2$ Trial | $0.971 \pm 0.005$ | $0.965 \pm 0.007$ | $0.965 \pm 0.006$ | $0.963 \pm 0.008$ | $0.976 \pm 0.005$ | $0.976 \pm 0.005$ |
| $R^2$ Average evoked | $0.976 \pm 0.010$ | $0.950 \pm 0.023$ | $0.961 \pm 0.016$ | $0.961 \pm 0.017$ | $0.972 \pm 0.013$ | $0.976 \pm 0.013$ |

**Fig 2:**
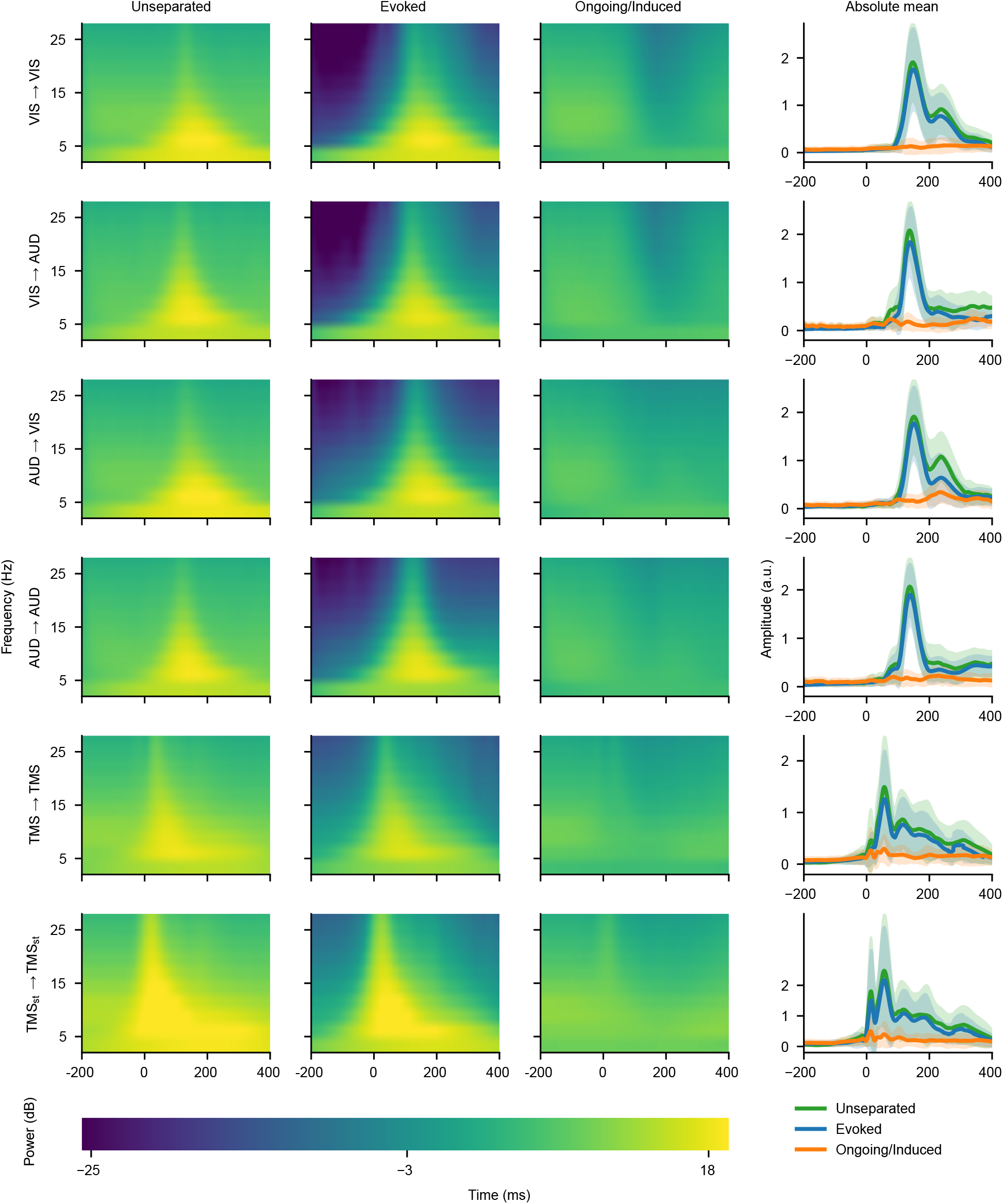
Mean power–frequency and absolute mean amplitude measures of recovered and unseparated (original data) response components across LOSO folds for all data modalities. The stimulation onset is at 0 ms. The standard deviation (SD) is depicted as shading of the respective component color. X*→*Y indicates that the model agnostic to the test subject was trained with data modality X and tested on modality Y.

### 2.2 Comparisons to alternative methods

For single-trial ground truth accuracy, we compared subject-agnostic single-trial decompositions obtained with EOIS to ablated variants, least-squares estimation (LSE) scaling, and dynamic time warping (DTW; Sakoe–Chiba window^17,18^ with 40-ms radius) of the mean evoked template computed excluding the test subject. This was done using intermixed pre-stimulus (ongoing) and modulated TMS-evoked EEG data as described in Section 5.1. EOIS outperformed all ablated variants—those lacking direct phase constraints, mean-evoked reconstruction, or single-trial reconstruction loss—as well as LSE and DTW-based approaches. Although LSE scaling and DTW can provide response estimates, their lower accuracies indicate limited separation accuracy and increased leakage between the components. The results are presented in Table 2.

**Table 2:** Single-trial response waveform accuracies (median *±* MAD of *R*^2^) for different subject-agnostic single-trial estimators from 0 to 500 ms of post-stimulus data. EOIS subscripts describe the ablated variants without specific losses (no phase: no PPC constraints, no mean: no average evoked reconstruction, and no single trial: not reconstructing the original data). LSE: Least-squares-scaled estimated waveform. DTW: Dynamic time-warped waveform (including LSE scaling after warping of z-scored waveforms). Significance stars indicate median differences from EOIS within metric (Holm-adjusted *p*-values from two-sided Wilcoxon signed-rank tests): ^*∗*^*p <* 10^*−*3^ and ^*∗∗*^*p <* 10^*−*8^.

| Method | Evoked | Ongoing |
| --- | --- | --- |
| EOIS | $0.61 \pm 0.14$ | $0.65 \pm 0.13$ |
| EOIS<sub>no phase</sub> | $-0.18 \pm 0.84$ ** | $0.02 \pm 0.01$ ** |
| EOIS<sub>no single trial</sub> | $0.53 \pm 0.16$ * | $-0.04 \pm 0.04$ ** |
| EOIS<sub>no mean</sub> | $-0.04 \pm 0.01$ ** | $0.27 \pm 0.42$ ** |
| LSE | $0.23 \pm 0.10$ ** | $0.34 \pm 0.39$ ** |
| DTW | $0.30 \pm 0.16$ ** | $0.46 \pm 0.17$ ** |

Across EOIS, LSE, and DTW, only the ongoing component recovery with EOIS was significantly correlated (*ρ ≈* 0.38, *p ≈* 0.02) with its respective signal-to-noise ratio (SNR); however, noise and component prevalence likely affect all components for all methods.

Mean-template subtraction and across-subject template morphing have limited utility for single-trial evoked–ongoing/induced separation because they directly assume similar timing and waveform shape across trials and subjects. Single-trial correlations (*ρ*) were moderate with within-subject templates and weaker with subject-agnostic templates, indicating low attainable fit quality (*ρ*^2^) for LSE scaling as shown in Table 3.

**Table 3:** Median *±* MAD of single-trial template–mixture fits (*ρ*^2^) of within-subject and subject-agnostic LSE-scaled mean evoked templates across subjects and trials. Because non-aligned components in the mixture attenuate template correlations, these values should be interpreted as conservative upper bounds on template–mixture fit quality rather than as evidence of evoked-component recovery.

| Modality | Visual | Auditory | TMS | TMS-st |
| --- | --- | --- | --- | --- |
| Within-subject | $0.50 \pm 0.11$ | $0.44 \pm 0.08$ | $0.37 \pm 0.21$ | $0.47 \pm 0.21$ |
| Subject-agnostic (LOSO) | $0.35 \pm 0.15$ | $0.28 \pm 0.11$ | $0.17 \pm 0.12$ | $0.25 \pm 0.16$ |

In the presence of non-Gaussian background activity, template morphing of weakly correlated templates is more likely to fit non-evoked structure, limiting its utility for single-trial separation of evoked and ongoing/induced components. Mean subtraction drives phase-locking in the residual toward zero by construction but enforces an across-trial constant evoked component and does not ensure a clean ongoing/induced component; amplitude differences, latency jitter, and morphology changes of the evoked response are pushed into the residual ongoing/induced component.

### 2.3 Pre-stimulus features bias post-stimulus inferences

We tested whether separating the components alters the estimated relationship between pre- and post-stimulus activity. Logistic regression was used to model single-trial high vs. low post-TMS activity based on pre-stimulus band-powers (theta, alpha, beta, and gamma) derived from within-trial pre–post causality-respecting pre-stimulus periods. Separate models were trained within subjects using the unseparated, evoked, and ongoing/induced components of the calibration data and the first 80% of application trials, and then tested on the last 20% of application trials. Post-stimulus magnitude was categorized by a median split of the root-mean-square (RMS) amplitudes within time windows (25–75, 75–200, and 25–300 ms), and performance was assessed using ROC–AUC.

Pre-stimulus features modeled post-stimulus non-phase-locked activity more effectively than evoked activity, with positive associations indicated by higher ROC–AUC values and odds ratios (Fig. 3). On average, the pre-stimulus RMS amplitudes were also more correlated with the post-stimulus ongoing/induced (*ρ ≈* 0.40) RMS amplitudes compared to the unseparated signal (*ρ ≈* 0.32) or evoked (*ρ ≈* 0.12) RMS amplitudes. These results demonstrate that associations obtained from the unseparated signal can largely reflect a modulated version of the non-phase-locked component rather than the evoked response.

**Fig 3:**
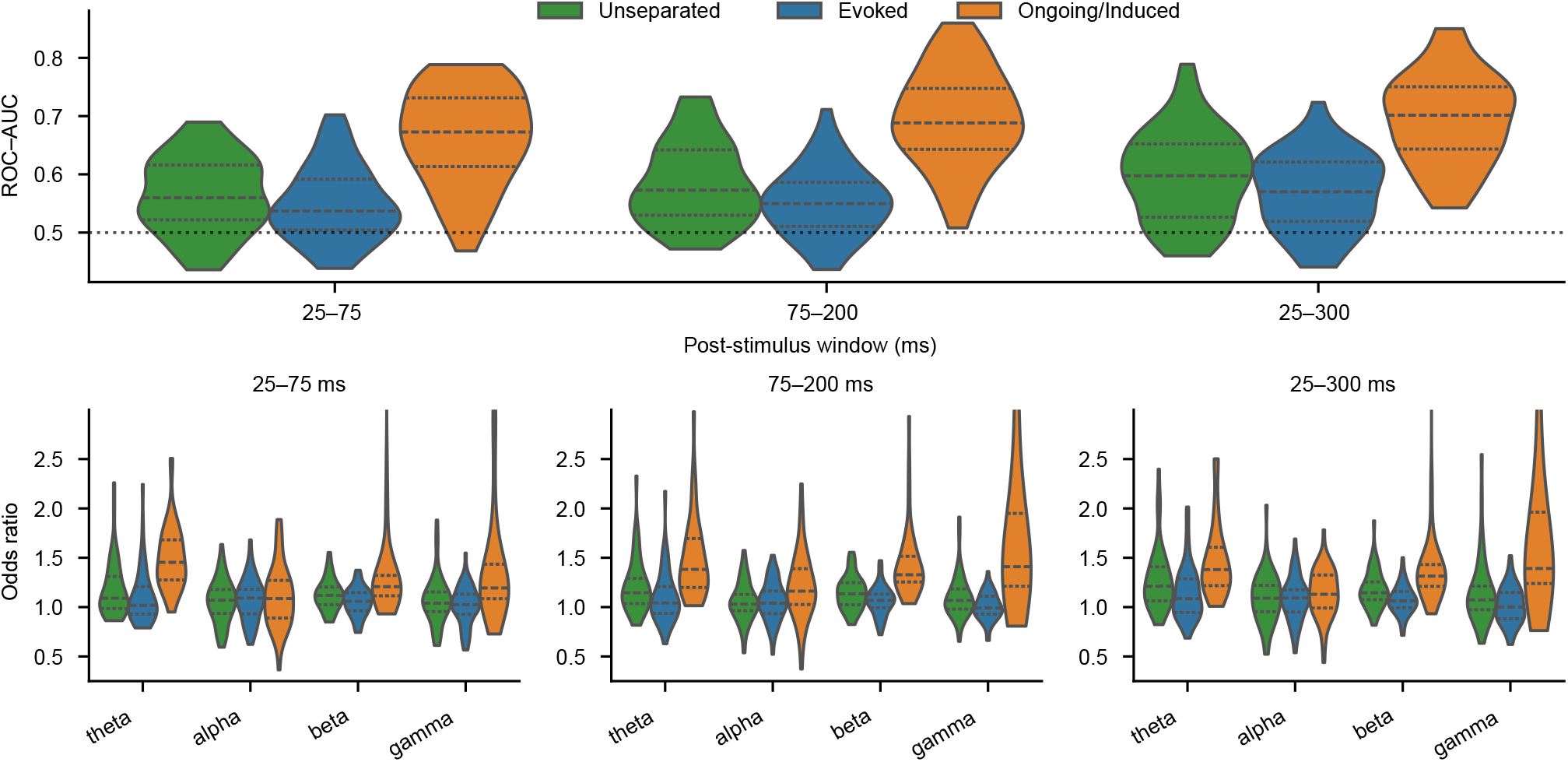
ROC–AUC and feature odds ratios across models for modeling the magnitude of post-stimulus components within defined time windows. Violin plots illustrate the across-subject distributions of ROC–AUC values and odds ratios for the frequency bands theta (4–7), alpha (8–12), beta (16–24), and gamma (30–40) Hz. Dotted lines within each violin plot indicate the distribution quartiles. For odds ratios, 1.0 represents the boundary, where values above indicate the feature increases the odds of a high-response outcome, and values below indicate a decreased likelihood. Note that although the input preprocessing windows are agnostic to the immediate post-stimulus responses, the post-stimulus-focused processing leverages pre-stimulus data for stability, which may inflate performance.

## 3 Discussion

Because EOIS operates on single-channel data, it allows flexible channel configurations. This streamlines EEG preparation for clinical use, facilitates analysis of previously recorded data, and supports real-time biomarkers by isolating evoked and ongoing/induced single-trial activity. EOIS also facilitates across-trial modeling of relationships between pre- and post-stimulus activity^1,5^ for both evoked and induced responses. Without component separation, post-stimulus evoked signals can reflect pre-stimulus non-phase-locked activity due to natural carry-over oscillations and temporal leakage, inflating performance and misleading inferences (Section 2.3). Induced activity can then be quantified as post-stimulus power in the non-phase-locked component relative to its pre-stimulus baseline.

The EOIS component decomposition takes 14 *±* 0.1 ms (median *±* MAD) for 1 s of 1-kHz data using a central processing unit. The calibrated and uncalibrated preprocessing pipelines enable real-time use with *<* 0.25 s processing per *≈* 1.6 s of 60-channel data. This makes EOIS especially useful for real-time closed-loop stimulation protocols^5,19^ and for TEP mapping^6,7^. In routine EEG without large stimulation-related artifacts, real-time preprocessing is already achievable^5,20^. Real-time monitoring and analysis of MEG data are also possible if the applied Maxwell filter and head-movement compensation^21,22^ operate causally or in short blocks. BCI applications may also benefit in performance and interpretability from separately processed evoked and induced responses. A key limitation of EOIS is phase-estimation uncertainty at short temporal scales. In M/EEG, sub-15-Hz rhythms carry most of the power, but their phases are difficult to estimate in short trials because too few cycles are present. In contrast, although higher-frequency activity typically contains enough cycles for phase estimation, low SNR can make these phases difficult to estimate. These uncertainties can cause leakage between evoked and ongoing/induced components. Although the implemented phase-consistency losses cannot fully control these uncertainties, average-reconstruction losses can still guide separation and reduce crosstalk. Naturally, longer trials should improve phase estimation and overall accuracy, but at the cost of greater computation and longer recording sessions. A further limitation is the lack of ground-truth labels for component separation at the single-trial level in real data: multiple single-trial decompositions can yield similar outcomes because labels are available only at the aggregate level. Our simulations for single-trial ground-truth evaluation also assume a purely additive model with modulated evoked activity.

Future work should prioritize pre-training on a broad range of response profiles drawn from multiple M/EEG stimulation modalities. Constructing a large pre-trained model and subsequently fine-tuning it for downstream tasks^5,9^ is likely to enhance cross-subject robustness and generalization across response types, while reducing the amount of fine-tuning data needed.

## 4 Conclusion

EOIS separates evoked and ongoing/induced activity at the single-trial level by leveraging phase-consistency features while remaining subject-agnostic. It supports offline M/EEG analysis as well as real-time EEG and TMS–EEG mapping to probe within- and across-trial dynamics. The framework yields separate per-trial waveform estimates, facilitating biomarker definition and deployment across individuals.

## 5 Methods

### 5.1 Dataset descriptions

We evaluated EOIS using real M/EEG datasets and semi-synthetic EEG derived from recorded data. The MEG data consisted of auditory- and visual-evoked fields from the CamCAN dataset^23^. The EEG data consisted of TMS-evoked potentials recorded in the ConnectToBrain project^1^. An overview of the protocols is given in Table 4.

**Table 4:** Dataset descriptions. A: The REFTEP dataset consists of recordings from both the University of Tübingen and Aalto University, where the channels and ITIs differ. M1: Primary motor cortex (left), rMT: Resting motor threshold^24^, ITI: Inter-trial interval, Mag and Grad: Magnetometers and gradiometers. *N*_subjects_: The number of subjects after exclusion criteria. The REFTEP data were subsampled to a common channel structure. For more information on the datasets, see CamCAN^23^ and REFTEP^1^.

| Name | CamCAN | REFTEP<sup>A</sup> |
| --- | --- | --- |
| Data type | MEG | EEG |
| Stimulation type | Visual & Auditory | Neuronavigated TMS |
| Stimulation protocol | Bilateral checkerboard & Binaural 300-Hz tones | M1 with $\gtrsim 110\%$ rMT<br>MagVenture/Nexstim |
| Blocks | 1 | 4 |
| Trials/Block | 60 Visual & 20 Auditory | 300 |
| Sampling rate (Hz) | 1000 | 5000 |
| ITI (s) | $1.0 \pm 0.2$ | $3.0 \pm 0.5$ & $4.25 \pm 0.25$ |
| Channels | 102 Mag + 204 Grad | 128 & 64 EEG |
| $N_{\text{subjects}}$ | 219 & 92 | 29 & 23 (52 total) |
| Male/Female | 129/90 & 56/36 | 24/28 |
| Age (years) | $54 \pm 18$ & $56 \pm 19$ | $28 \pm 6$ |

For ground truth evaluations, we generated a synthetic TMS-evoked EEG dataset by mixing 1 s of pre-stimulus data with a semi-synthetic evoked signal for each trial from data preprocessed as in section 5.3.1. The evoked signal was constructed, for each subject, as a randomly modulated version of that subject’s average evoked waveform. At most 650 trials per subject were included for training and testing on simulated data. To ensure a robust evoked-response estimate, we included only subjects (*N* = 36) whose averaged calibration period signal met the following criteria: (*i*) the channel C3 had a peak-to-peak amplitude of *≥* 5 *µ*V during 20 to 150 ms; and (*ii*) the RMS value of the post-stimulus window from 20 to 150 ms was at least 10 times the RMS value of the pre-stimulus window from −500 to −100 ms.

Amplitude modulation was applied in two stages. First, we created a smooth, time-varying envelope over the evoked segment by sampling Gaussian noise at each time point from *N* (0, 1), smoothing it with a Gaussian kernel with an SD of 40 ms, and rescaling it to achieve a target SD of 0.5. This modulation was then converted to a strictly positive, multiplicative envelope using a lognormal transform. Second, a global, trial-specific gain was drawn from *N* (1, 0.5) and clipped to a minimum of 0.3. The evoked component for each trial was the subject-specific template multiplied by this envelope and gain, and the synthetic trial was obtained by adding the evoked component to the pre-stimulus data.

### 5.2 Visual and auditory MEG

The MaxFilter™ 2.2.15 (Elekta Neuromag) was applied for noise suppression, head movement compensation, and bad channel detection and reconstruction. The data were band-pass filtered between 1 and 30 Hz (Hamming-windowed finite impulse response). Independent component analysis (ICA)^25,26^ was used to identify and suppress ocular and cardiac artifacts based on outlying absolute correlations (z-score *>* 3) with the electrooculogram (EOG) and electrocardiogram channels. Continuous MEG data were epoched from −200 to 700 ms relative to event onset. Trials were rejected if their SD z-score exceeded 2.5 or if the absolute amplitude in either EOG channel exceeded a z-score threshold of 3 during −100 to 100 ms. The data were baseline corrected using the mean from −200 to 0 ms.

Because the number of trials was small (*≤* 60 or *≤* 20 for visual and auditory stimuli, respectively), we applied an inclusion criterion to ensure robust average evoked-response estimates, under which ongoing/induced activity and noise were expected to average to near zero across trials relative to the evoked response. We included only subjects for whom the ratio between the RMS value of the averaged post-stimulus period from 100 to 300 ms and the pre-stimulus period from −200 to 0 ms exceeded 15 (visual) or 10 (auditory). For each subject and condition, we selected a representative channel with the largest aforementioned RMS. The epoched data were normalized by the total RMS value and adjusted to positive polarity using the sign of the area under the curve (AUC) from 100 to 200 ms from the average response. Note that while this specific channel-selection method cannot be performed online, in practice, the experimenter can select a channel in real time, and EOIS can be applied to both polarities. The IC decompositions and rejection criteria can also be calibrated from previously recorded resting state data to support across-trial causality.

### 5.3 TMS–EEG

#### 5.3.1 Real-time processing with an offline calibration phase

TMS-evoked EEG data were preprocessed using an offline calibration phase followed by a simulated real-time application phase^5,10^.

The calibration phase consisted of 200 trials. Using pre- and post-stimulus periods, we separately calibrated individual filters: z-score-based channel rejection (pre: lag-1 autocorrelation |*z*| *<* 4 and MAD *z ∈/* [*−*6, 4]; post: AUC *z >* 2.5) and channel interpolation matrices (spherical spline). Trial rejection used z-score distributions based on MAD (pre: *z ∈/* [*−*8, 5]; post: *z >* 6), and trials were also rejected if a single channel MAD had *z >* 5 relative to other channels. The rejection criterion was iterated until no additional bad trials or channels were rejected. We applied ocular IC filters derived from pre-stimulus data^25,27^ using ICLabel^28^, and fixed-component post-stimulus-calibrated muscle-artifact-suppressing signal-space projection and source-informed reconstruction (SSP–SIR) filters^5,29,30^ calibrated from post-stimulus data. The calibrated filters were subsequently applied to single trials in the application phase.

The data were filtered to 2–40 Hz (zero-phase 2nd order Butterworth) within trials from −1100 to 600 ms. Pulse artifacts were initially interpolated using a piecewise cubic Hermite interpolating polynomial (PCHIP) between −4 and 6 ms and re-interpolated between −6 and 15 ms after SSP–SIR. The data were polyphase-downsampled to 1 kHz, mean baseline-corrected using data from −500 to −25 ms, and then re-average referenced. Channel C3 near the stimulation site was used for TEP and pre-stimulus readouts. The post-stimulus data were cropped from −500 to 500 ms and normalized by the RMS value calculated from the calibration data. For causal inference purposes, the pre-stimulus periods were additionally processed separately from the post-stimulus-focused data using the aforementioned trial rejection and channel recovery criteria, ICA, and filtering.

#### 5.3.2 Real-time processing without calibrated filters

For real-time preprocessing without prior calibration, we used filters derived only from the current trial, −1000 to 600 ms relative to TMS, to mitigate the effect of biasing spatially-dependent TMS-related artifacts.

The data were baseline corrected using the mean from −500 to −25 ms. We then performed PCHIP interpolation for the TMS pulse artifact from −4 to 6 ms. Bad channels and trials were identified and either interpolated (spherical spline) or rejected using within-trial statistics computed separately for pre- and post-stimulus periods: pre-stimulus metrics include across-channel z-scored lag-1 autocorrelations (*z ∈/* [−3, 10]) and SDs (*z ∈/* [−10, 3]), and the post-stimulus metric from 20 to 50 ms is the across-channel z-scored AUC (*z >* 3.5). The rejection procedure was iterated until no additional channels were rejected; trials with more than 15% rejected channels were excluded. An SSP–SIR filter^5,29,30^ was calibrated from 7 to 35 ms and applied to suppress TMS-evoked muscle artifacts. An extended recovered pulse artifact window from −5 to 10 ms was then interpolated using PCHIP, the data were filtered to 2–40 Hz (zero-phase 2nd order Butterworth), polyphase-downsampled to 1 kHz, re-baseline corrected and average referenced, and cropped from −500 to 500 ms. After preprocessing, the data were normalized by the RMS value calculated from the first valid trial for causal across-trial comparisons and numeric stability.

### 5.4 Model architecture

EOIS decomposes individual trials **x**_*i*_ into phase-locked evoked **e**_*i*_ and non-phase-locked ongoing/induced components **u**_*i*_, such that **x**_*i*_ = **e**_*i*_ + **u**_*i*_, without imposing cross-component independence. The model primarily enforces three constraints: (*i*) the across-trial mean of the evoked components should converge to the across-trial mean of the original data, because non-phase-locked components in the data converge to zero in expectation; (*ii*) because the components are estimated using separate heads, the mean ongoing/induced component should converge to zero across trials; and (*iii*) the evoked and ongoing/induced components should show high and low phase consistency across trials, respectively. The model architecture is presented in Fig. 4 and was implemented in PyTorch^31^.

**Fig 4:**
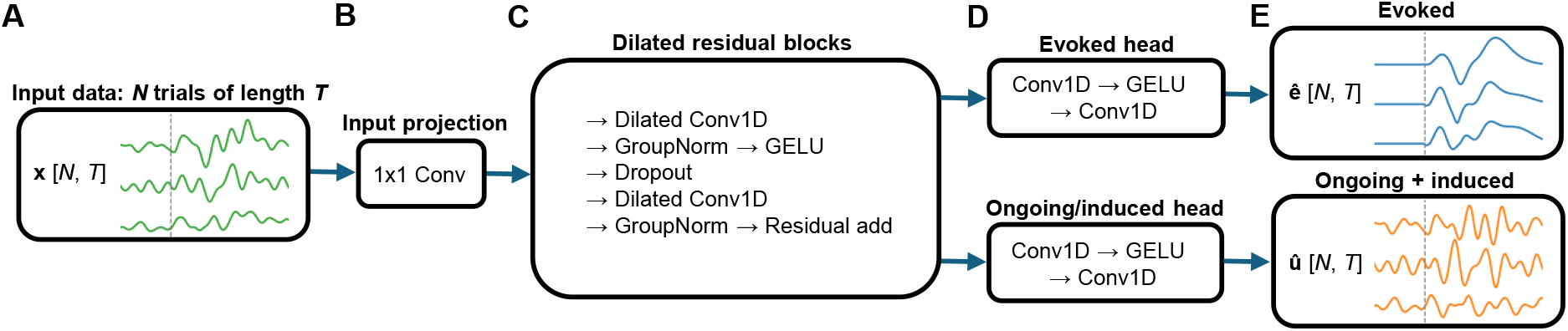
EOIS model architecture. **A**: Each trial in the input data includes pre- and post-stimulus activity, where the dashed line depicts the event onset. **B**: An input 1×1 convolution projects the signal to a 64-channel latent space. **C**: Residual blocks use stacked dilated 1D-convolutions^12–15^ (kernel sizes 5, 5, 7, 9, 9, and 9 for respective dilations 1, 2, 4, 8, 16, and 32) to identify temporal context at multiple scales, group normalization (GroupNorm), Gaussian error linear unit (GELU), and a dropout (*p* = 0.1). **D**: The shared stack feeds the evoked and ongoing/induced heads resulting in the respective component estimates (**E**). The evoked head convolutions have a kernel size of 3 for encouraging consistent within-response temporal structure, while the ongoing/induced head convolutions have a kernel size of 1.

#### 5.4.1 Training

We used LOSO cross-validation: in each outer fold, one subject was held out for testing, and the remaining subjects were randomly split into training (70%) and validation (30%). The model with the lowest validation loss was selected. A summary of the training procedure is presented in Table 5.

**Table 5:** Training configuration and evaluation protocol. Each batch consists of trials from a single subject. LR: learning rate. RoP: Reduce on plateau. The validation loss was smoothed using the mean over the previous 5 epochs.

| Component | Setting |
| --- | --- |
| Loss function | Custom, see Section 5.4.2 |
| Optimizer | AdamW |
| Scheduler | Reduce on plateau (RoP) |
| RoP settings | threshold = $10^{-4}$, factor = 0.5, patience = 10 epochs |
| RoP maximum LR | $5 \cdot 10^{-4}$ |
| RoP minimum LR | $10^{-7}$ |
| Warmup | 10 epochs |
| Weight decay | $10^{-5}$ |
| Batch size | min(trials of subject, 200) |
| Early stopping | $\Delta_{\min} = 10^{-4}$, patience = 30 epochs |
| Max. train epochs | 300 |

#### 5.4.2 Loss function

We use known convergent features of phase-locked and non-phase-loscked activity across a high number of within-subject trials to define extraction and reconstruction losses. The training is guided by the minimization of the sum of the feature losses *L*_f_, described in the following paragraphs, as min ∑_f_ *λ*_f_*L*_f_, where *λ*_f_ is the specific feature weight.

#### Single-trial reconstruction

EOIS learns phase-locked and non-phase-locked activity separately. The estimated phase-locked component **ê** and non-phase-lockedûcomponents should reconstruct the original data **x** in each trial. We therefore define the reconstruction loss across time and trials as the normalized mean squared error (MSE) with *λ*_reco_ = 3.0 as

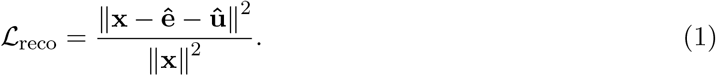

With sufficiently long and jittered ITIs, pre-stimulus activity is expected to contain minimal stimulus-related phase-locked components. We therefore letûwith the loss dominate the pre-stimulus data with the loss

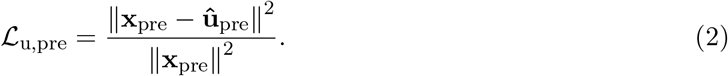

Because this loss operates sample-by-sample and has little competition, we use *λ*_u,pre_ = 1.0.

#### Average reconstruction

The across-trial average of non-phase-locked ongoing/induced activity and non-event-related noise converges to zero in time. As a result, the across-trial average of the phase-locked evoked components converges to the across-trial average of the individual trials **x**. We therefore define the evoked response reconstruction loss as the strongly enforced (*λ*_e,post_ = 8.0) normalized MSE during the post-stimulus times as

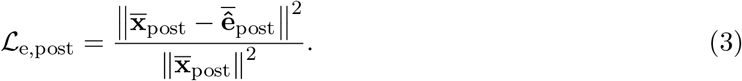

Similarly, we enforce that the across-trial average of the non-phase-locked components converges to zero across trials using the normalized MSE from zero, with *λ*_u,post_ = 3.0 as

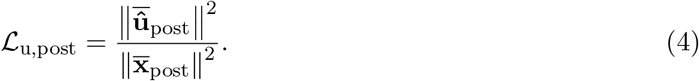

#### Phase coherence

Evoked activity is phase-locked to the stimulus and therefore expected to show high across-trial phase consistency. By contrast, non-phase-locked activity is expected to show low across-trial phase consistency. However, evoked components can jitter in timing and amplitude^1^, which can be unfairly penalized by strict phase-locking measures in nonstationary and low SNR conditions. The ground truth phase consistency of evoked components across trials is therefore expected to be high, but below one. We encourage the evoked component to have high phase-locking by minimizing a focal, rectified (non-zero ReLU) pairwise phase consistency (PPC_R_)^16^ loss computed from its Morlet transform **Ê** during post-stimulus times as

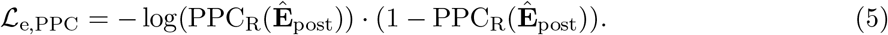

In contrast, the non-phase-locked component is guided to have minimal phase consistency computed from its Morlet transform Û by minimizing the PPC as

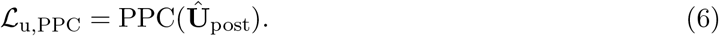

Because evoked consistency is already enforced by the average reconstruction, and to avoid artificial phase-locking, we use *λ*_e,PPC_ = 3.0, while the ongoing/induced component is driven to zero phase-locking with *λ*_u,PPC_ = 6.0. To balance time–frequency resolutions in Morlet wavelet–transformed data used to calculate the PPCs for frequencies over 4 Hz, we used a piecewise cycle schedule: 3 for *f <* 6, 4 for 6 *≤ f <* 12, 5 for 12 *≤ f <* 20, and 6 for *f ≥* 20 Hz.

#### Canceling artifacts

Because EOIS learns evoked and ongoing/induced components separately and enforces that they sum to the original data, it is possible that both components contain artifacts of opposite sign. We therefore encourage the sum of the component powers not to exceed the power of the data in padded, within-window-averaged trials, denoted by (*·*)_*w*_ (window length 20% of the trial; hop 10%) with the loss (*λ*_power_ = 1.0)

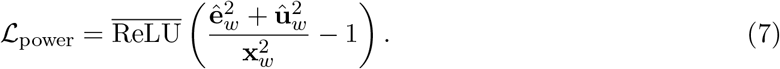

#### 5.4.3 Evaluation and statistical analysis

We evaluated the results using coefficients of determination of the evoked and ongoing/induced activity waveforms during post-stimulus times. Specifically, for the real M/EEG data, we evaluated the single-trial reconstruction accuracy and the recovery of the averaged evoked component. In synthetic data, where ground-truth labels are available, we separately evaluated the accuracy of both components across trials. The across-trial phase consistencies of the components were calculated using pairwise phase consistency^16^.

## Acknowledgements

This work was funded by the Jenny and Antti Wihuri Foundation (Grant No. 00240097).

The REFTEP data were collected within the ConnectToBrain project funded by the European Research Council (ERC) under the European Union’s Horizon 2020 research and innovation programme (Grant No. 810377).

CamCAN funding was provided by the UK Biotechnology and Biological Sciences Research Council (Grant No. BB/H008217/1), together with support from the United Kingdom Medical Research Council and University of Cambridge, United Kingdom.

We acknowledge the use of computational resources provided by the Triton computing cluster of the Aalto Science-IT project. The OpenAI GPT-based Aalto AI Assistant provided by Aalto University was used for architectural design and code development assistance.

## Data availability

De-identified EEG data are available from the corresponding authors upon reasonable request, subject to ethical approval and data agreements. The MEG data were obtained from the CamCAN repository^23^ (available at https://cam-can.mrc-cbu.cam.ac.uk/dataset/).

## Code availability

The code used in this study is publicly available and can be accessed at https://github.com/OAhola/EOIS.

## References

[1] Ahola, O. et al. “Predictive modeling of TMS-evoked responses: Unraveling instantaneous excitability states”. In: NeuroImage 322 (2025). doi: 10.1016/j.neuroimage.2025.121553.

[2] Haxel, L. et al. “Decoding motor excitability in TMS using EEG-features: an exploratory machine learning approach”. In: IEEE Transactions on Neural Systems and Rehabilitation Engineering 33 (2025). doi: 10.1109/TNSRE.2024.3516393.

[3] Zrenner, C., Desideri, D., Belardinelli, P., and Ziemann, U. “Real-time EEG-defined excitability states determine efficacy of TMS-induced plasticity in human motor cortex”. In: Brain Stimulation 11.2 (2018). doi: 10.1016/j.brs.2017.11.016.

[4] Van Dijk, H., Schoffelen, J.-M., Oostenveld, R., and Jensen, O. “Prestimulus oscillatory activity in the alpha band predicts visual discrimination ability”. In: The Journal of Neuroscience 28.8 (2008). doi: 10.1523/JNEUROSCI.1853-07.2008.

[5] Haxel, L., Ahola, O., Kapoor, J., Ziemann, U., and Macke, J. H. “Personalized real-time inference of momentary excitability from human EEG”. In: NeuroImage 322 (2025). doi: 10.1016/j.neuroimage.2025.121547.

[6] Casarotto, S. et al. “The rt-TEP tool: real-time visualization of TMS-evoked potentials to maximize cortical activation and minimize artifacts”. In: Journal of Neuroscience Methods 370 (2022). doi: 10.1016/j.jneumeth.2022.109486.

[7] Ukharova, E. et al. “Neuroimaging-guided TMS–EEG for real-time cortical network mapping”. In: JoVE 220 (2025). doi: 10.3791/67339.

[8] Blankertz, B., Lemm, S., Treder, M., Haufe, S., and Müller, K.-R. “Single-trial analysis and classification of ERP components — a tutorial”. In: NeuroImage 56.2 (2011). Multivariate Decoding and Brain Reading. doi: 10.1016/j.neuroimage.2010.06.048.

[9] Haxel, L., Kapoor, J., Ziemann, U., and Macke, J. H. “EDAPT: Towards calibration-free BCIs with continual online adaptation”. In: Journal of Neural Engineering 23.2 (2026). doi: 10.1088/1741-2552/ae5689.

[10] Ahola, O., Haxel, L., and Ziemann, U. “Automated spatiotemporal response identification and separation for averaged and single-trial EEG and MEG data”. In: IEEE Transactions on Biomedical Engineering (2026). doi: 10.1109/TBME.2026.3676122.

[11] Jung, T.-P. et al. “Analysis and visualization of single-trial event-related potentials”. In: Human Brain Mapping 14.3 (2001). doi: 10.1002/hbm.1050.

[12] He, K., Zhang, X., Ren, S., and Sun, J. “Deep residual learning for image recognition”. In: Proceedings of the IEEE Conference on Computer Vision and Pattern Recognition. 2016. doi: 10.1109/CVPR.2016.90.

[13] Bai, S., Kolter, J. Z., and Koltun, V. “An empirical evaluation of generic convolutional and recurrent networks for sequence modeling”. In: arXiv (2018). doi: 10.48550/arXiv.1803.01271.

[14] Yu, F., Koltun, V., and Funkhouser, T. “Dilated residual networks”. In: Proceedings of the IEEE Conference on Computer Vision and Pattern Recognition. 2017. doi: 10.1109/CVPR.2017.75.

[15] van den Oord, A. et al. “WaveNet: a generative model for raw audio”. In: arXiv (2016). doi: 10.48550/arXiv.1609.03499.

[16] Vinck, M., van Wingerden, M., Womelsdorf, T., Fries, P., and Pennartz, C. M. “The pairwise phase consistency: a bias-free measure of rhythmic neuronal synchronization”. In: NeuroImage 51.1 (2010). doi: 10.1016/j.neuroimage.2010.01.073.

[17] Sakoe, H. and Chiba, S. “Dynamic programming algorithm optimization for spoken word recognition”. In: IEEE Transactions on Acoustics, Speech, and Signal Processing 26.1 (1978). doi: 10.1109/TASSP.1978.1163055.

[18] Tavenard, R. et al. “Tslearn, a machine learning toolkit for time series data”. In: Journal of Machine Learning Research 21.118 (2020). doi: 10.5555/3455716.3455834.

[19] Kahilakoski, O.-P. et al. “NeuroSimo: an open-source software for closed-loop EEG-or EMG-guided TMS”. In: bioRxiv (2025). doi: 10.1101/2025.04.05.647342.

[20] Schalk, G., McFarland, D. J., Hinterberger, T., Birbaumer, N., and Wolpaw, J. R. “BCI2000: a general-purpose brain-computer interface (BCI) system”. In: IEEE Transactions on Biomedical Engineering 51.6 (2004). doi: 10.1109/TBME.2004.827072.

[21] Taulu, S. and Kajola, M. “Presentation of electromagnetic multichannel data: The signal space separation method”. In: Journal of Applied Physics 97.12 (2005). doi: 10.1063/1.1935742.

[22] Taulu, S. and Simola, J. “Spatiotemporal signal space separation method for rejecting nearby interference in MEG measurements”. In: Physics in Medicine and Biology 51.7 (2006). doi: 10.1088/0031-9155/51/7/008.

[23] Shafto, M. A. et al. “The Cambridge Centre for Ageing and Neuroscience (Cam-CAN) study protocol: a cross-sectional, lifespan, multidisciplinary examination of healthy cognitive ageing”. In: BMC Neurology 14 (2014). doi: 10.1186/s12883-014-0204-1.

[24] Rossini, P. M. et al. “Non-invasive electrical and magnetic stimulation of the brain, spinal cord, roots and peripheral nerves: Basic principles and procedures for routine clinical and research application. An updated report from an I.F.C.N. Committee”. In: Clinical Neurophysiology: Official Journal of the International Federation of Clinical Neurophysiology 126.6 (2015). doi: 10.1016/j.clinph.2015.02.001.

[25] Gramfort, A. et al. “MEG and EEG data analysis with MNE-Python”. In: Frontiers in Neuroscience 7 (2013). doi: 10.3389/fnins.2013.00267.

[26] Hyvärinen, A. “Fast and robust fixed-point algorithms for independent component analysis”. In: IEEE Transactions on Neural Networks 10.3 (1999). doi: 10.1109/72.761722.

[27] Lee, T.-W., Girolami, M., and Sejnowski, T. J. “Independent component analysis using an extended infomax algorithm for mixed subgaussian and supergaussian sources”. In: Neural Computation 11.2 (1999). doi: 10.1162/089976699300016719.

[28] Pion-Tonachini, L., Kreutz-Delgado, K., and Makeig, S. “ICLabel: An automated electroencephalographic independent component classifier, dataset, and website”. In: NeuroImage 198 (2019). doi: 10.1016/j.neuroimage.2019.05.026.

[29] Mutanen, T. P. et al. “Recovering TMS-evoked EEG responses masked by muscle artifacts”. In: NeuroImage 139 (2016). doi: 10.1016/j.neuroimage.2016.05.028.

[30] Rogasch, N. C. et al. “Analysing concurrent transcranial magnetic stimulation and electroencephalographic data: A review and introduction to the open-source TESA software”. In: NeuroImage 147 (2017). doi: 10.1016/j.neuroimage.2016.10.031.

[31] Paszke, A. et al. “PyTorch: an imperative style, high-performance deep learning library”. In: arXiv (2019). doi: 10.48550/arXiv.1912.01703.

